# Highly efficient and versatile plasmid-based gene editing in primary T cells

**DOI:** 10.1101/247544

**Authors:** Mara Kornete, Romina Marone, Lukas T. Jeker

## Abstract

Adoptive cell transfer (ACT) is an important approach for basic research and emerges as an effective treatment for various diseases including infections and blood cancers. Direct genetic manipulation of primary immune cells opens up unprecedented research opportunities and could be applied to enhance cellular therapeutic products. Here, we report highly efficient genome engineering in primary murine T cells using a plasmid-based RNA-guided CRISPR system. We developed a straightforward approach to ablate genes in up to 90% of cells and to introduce precisely targeted single nucleotide polymorphisms (SNP) in up to 25% of the transfected primary T cells. We used gene editing-mediated allele switching to quantify homology directed repair (HDR), systematically optimize experimental parameters and map a native B cell epitope in primary T cells. Allele switching of a surrogate cell surface marker can be used to enrich cells with successful simultaneous editing of a second gene of interest. Finally, we applied the approach to correct two disease-causing mutations in the *Foxp3* gene. Both repairing the cause of the scurfy syndrome, a 2bp insertion in *Foxp3*, and repairing the clinically relevant Foxp3^K276X^ mutation restored Foxp3 expression in primary T cells.

## Introduction

Lymphocytes are among the best understood mammalian cells. A plethora of genetically modified mice have yielded deep insight into the molecular and cellular processes underlying lymphocyte, but also more generally, mammalian development and function. Inbred mouse strains enable adoptive transfer experiments without immunological rejection. However, although genetically modified murine models are very powerful, a major practical limitation is the time required to generate genetically altered mice. Moreover, intercrossing mice with combinations of mutations and/or transgenes requires extensive breeding. Finally, for immunologic reasons a given genetic alteration often either needs to be introduced on a particular genetic background, or mice need to be backcrossed for multiple generations to change the genetic background. Thus, systems to directly genetically edit murine lymphocytes would be highly desirable to reduce the need for breeding.

While previous efforts to establish CRISPR/Cas9-mediated gene editing in primary T cells have mainly focused on human T cells (1), (2), (3), (4) to our knowledge only two reports describe successful CRISPR/Cas9 gene editing in primary murine T cells (5), (6). Both approaches depend on mice expressing transgenic Cas9 and either a second transgenic construct to express the gRNA (5), or viral transduction of T cells followed by antibiotic selection (6). As a result, although these approaches constitute significant advances, they still require breeding of a constitutive Cas9 transgene, which may be genotoxic and might lead to immunologic rejection after adoptive transfer of transgenic cells.

In an earlier report transfection of plasmids for CRISPR/Cas9-mediated genome editing in human hematopoietic stem cells (hHSC) and CD4^+^ T cells was successfully used (1). However, gene ablation was less efficient in T cells than hHSCs with efficiencies in primary human CD4^+^ T cells mostly <10%, even with a strategy using two gRNAs targeting the same gene simultaneously. Subsequently, it was reported that CRISPR/Cas9 genome engineering could be improved in primary human T cells by replacing plasmids with chemically modified guide RNAs combined with Cas9 encoding mRNA (3). Nucleofection with a plasmid encoding both the sgRNA and Cas9 did not demonstrate editing efficiencies above background while co-transfection of a chemically modified sgRNA with a Cas9 expressing plasmid yielded deletion in <10% of T cells. This is in comparison to high double digit deletion efficiencies with sgRNA and Cas9 mRNA (3). Similarly, electroporation of recombinant Cas9/gRNA ribonucleoprotein complexes (RNPs) results in moderate to high double digit deletion efficiencies (2). More recently, multiplexed high efficiency CRISPR/Cas9 editing was reported using mRNA and multiple in vitro transcribed sgRNAs (4). Thus, there is a growing notion that in comparison to other approaches DNA-based techniques work poorly, if at all, for CRISPR/Cas9 gene editing in primary T cells (2), (3), (7), (8). However, as plasmids have been the workhorse of molecular biology for decades because of their ease of use, versatility, low production costs and their wide availability to the scientific community they would have specific merits were they to be successfully used. Importantly, thousands of CRISPR-related products are already available as plasmids and the latest CRISPR nuclease variants are rapidly being deposited to a growing, publically available resource (Addgene.org/crispr). In addition, in contrast to viral transduction and/or transgenic Cas9 expression, transient expression of nuclease and gRNA is sufficient and even preferred as a method to reduce off-target genome editing. Furthermore, technical advances make electroporation a clinically useful approach (9). Finally, RNPs are more difficult to produce than plasmids, particularly for complex constructs such as base editors, and are more expensive than plasmids thus preventing access to low budget research groups.

We report here a highly efficient method to ablate genes in primary murine T cells. The approach is based on plasmids that are commonly accessible through publically available resources (Addgene.org). Combining two plasmids enables efficient multiplexed gene ablation. Gene edited T cells are viable, home to lymphoid organs, expand in a lymphophenic environment and differentiate normally in response to a viral infection in lymphoreplete mice. Using a novel assay, we additionally increased the efficiency of HDR in primary CD4^+^ T cells reaching up to 30% HDR using commonly available reagents. Through the application of this approach we mapped a native B cell epitope directly in primary cells and corrected a defective Foxp3 gene. We demonstrate that multiplexed HDR editing of a cell surface molecule can serve as a surrogate marker to enrich for cells undergoing HDR at a second locus for which no marker may exist, similar to previous findings (10). This principle was applied to enrich gene-corrected Foxp3-deficient cells. Thus, we report a versatile T cell editing approach, and provide proof-of-principle that gene-edited Foxp3-deficient T cells can produce the repaired gene product, Foxp3 protein.

## Materials and Methods

### Gene editing in primary murine CD4^+^ T cells

Naïve CD4^+^ T cells were purified (>96% purity) from C57BL/6N mouse spleen (SP) and lymph nodes (LN) using the EasySep^TM^ Mouse Naïve CD4^+^ T Cell Isolation Kit (STEMCELL Technologies Inc). Alternatively, total CD4^+^ T cells were purified (>96% purity) from C57BL/6 SMARTA^+^ CD45.1 mouse SP and LN using the EasySep^TM^ Mouse CD4^+^ T Cell Isolation Kit (STEMCELL Technologies Inc). Complete RPMI media (CM RPMI) was generated by supplementing RPMI (Sigma) with 10% heat-inactivated FCS (Atlanta biologicals), 2mM Glutamax (Gibco), 50µM β-mercaptoethanol (Gibco), 10mM HEPES (Sigma) and non essential amino acids (Gibco). For T cell activation, 2×10^6^ naïve CD4^+^ T cells were plated in a 24-well plate (Corning) coated with monoclonal antibodies (mAb) anti-CD3 (hybridoma clone 2C11, 1µg/ml) and anti-CD28 (hybridoma clone PV-1, 0.5µg/ml, both BioXcell) for 24h at 37°C with 5% CO2 in the presence of 50IU/ml recombinant human Interleukin-2 (rhIL-2) (RD systems). 24h later T cells were harvested and washed with PBS. 2×10^6^ activated T cells were electroporated with the Invitrogen Neon® Transfection System (Invitrogen) at the following conditions: voltage (1550V), width (10ms), pulses (3), 100µl tip, buffer R. Cells were transfected with 6.5µg of empty plasmid px458 (Addgene plasmid number 48138) or the plasmids described in Figure legends and Table S1. (Addgene plasmid numbers 82670, 82672-82675, 82677, 102910). For HDR cells were co-transfected with 12µg HDR template (if plasmid: Table S3; Addgene 82661-82665, 82667, 104988) or 10µl of 10µM stock ssDNA template (Table S) purchased from IDT. After electroporation cells were plated in 24-well plate in 650µl CM RPMI with 50IU rhIL-2/ml in the presence of plate-bound mAbs at half the concentrations used for the initial activation, i.e. anti-CD3 (0.5µg/ml) and anti-CD28 (0.25µg/ml). GFP^+^ and GFP- cells were sorted 24h post transfection using a FACSAria Cell Sorter to >98% purity (BD Biosciences). Immediately after sorting cells were plated in 96 well flat bottom plates without activating antibodies in 250µl CM RPMI supplemented with 50IU rhIL-2/ml. For the HDR experiments sorted cells were cultured in the presence of NHEJ inhibitors or HDR enhancers for the following 24h in order to enhance HDR (as indicated in figure legends). Cells were re-activated with plate bound anti-CD3 (0.5µg/ml) and anti-CD28 (0.25µg/ml) on day 4 post GFP sorting and expanded for the following 9 days in culture until the end of the experiment. For in vivo editing experiment cells were injected immediately into the recipient mouse.

### Gene editing in EL-4 cells

EL-4 cells were grown in RPMI (Sigma) supplemented with 10% heat inactivated fetal bovine serum (Atlanta biologicals), 2mM Glutamax (Gibco) and 50µM β-mercaptoethanol (Gibco). FACS analysis confirmed homozygous CD90.2 and CD45.2 expression by EL-4 cells comparable to that of primary T cells. 2×10^6^ EL-4 cells were electroporated with the Invitrogen Neon® Transfection System (Invitrogen) at the following conditions: voltage (1080V), width (50ms), number of pulses (1), 100µl tip. The amount of plasmids and concentrations of HDR templates were the same as for the primary T cells described above. After electroporation cells were plated in 24 well plates in 650µl CM RPMI. GFP^+^ and GFP- cells were sorted 24h post transfection using a FACSAria Cell Sorter (BD Biosciences) to a purity of >98%. Immediately after sorting cells were plated in 96 well flat bottom plates. For the HDR experiments, sorted cells were cultured in the presence of NHEJ inhibitors or HDR enhancers for the following 24h in order to enhance the HDR. Cells were then expanded for the next 7-9 days in culture.

### Foxp3 repair protocol

Although the majority of T cells from Foxp3^K276X^ C57BL/6 mice are highly activated, they had to be re-activated in vitro for electroporation, otherwise we could not obtain reasonable transfection efficiencies. We adjusted the protocol used to electroporate primary T cells from healthy mice by reducing the TCR stimulation in order to obtain a good balance between cell viability and transfection rate. In addition, we used total CD4^+^ T cells as a starting population because of the low numbers of naïve T cells (data not shown). Total CD4^+^ T cells were purified from Foxp3^K276X^ C57BL/6 mice pooled SP and LN using the EasySep^TM^ CD4^+^ T Cell Isolation Kit (>96% purity) (STEMCELL Technologies Inc). For T cell activation, 2×10^6^ CD4^+^ T cells were plated in a 24-well plate coated with anti-CD3 (clone 2C11; 0.5µg/ml) and anti-CD28 (clone PV-1; 0.25µg/ml) (BioXcell) for 24h at 37 °C with 5% CO2, with 50IU rhIL- 2/ml (RD systems). 24h later T cells were harvested and washed with PBS. 2×10^6^ activated T cells were electroporated with the Invitrogen Neon® Transfection System (Invitrogen) at the following conditions: voltage (1550V), width (10ms), number of pulses (3). Cells were transfected with 6.5µg of plasmid (p240_LTJ_sgRNAFoxp3^K276X^, Addgene number 82675) and 12µg of the dsDNA wt Foxp3 repair template (Addgene 82664). After electroporation cells were plated in 24 well plate with 50IU/ml of rhIL-2 in the presence of plate bound mAbs at half the concentrations used for the initial activation, i.e. 0.25µg/ml anti-CD3 and 0.12µg/ml anti-CD28 in 650µl CM RPMI. GFP^+^ and GFP^−^ cells were sorted 24h post transfection using a FACSAria Cell Sorter to a purity >98% (BD Biosciences).

Immediately after cell sorting the purified cells were re-activated with plate bound anti-CD3 (0.5µg/ml) and anti-CD28 (0.25µg/ml) and expanded until the end of the experiment in the presence of rhIL-2 (250IU/ml), TGFβ (5ng/ml, RD Systems), anti- IFNγ (10mg/ml, BioXcell), anti-IL-4 (10mg/ml, BioXcell) and with or without Retinoic Acid (RA) (10mM, Sigma) as indicated in the figure legend.

### Mice

C57BL/6N (Charles River stock No: 027) were purchased at the Charles River laboratory. Foxp3^K276X^ C57BL/6 (Jackson laboratory Stock No: 019933) mice were a generous gift from Ed Palmer (Basel University Hospital). C57BL/6 SMARTA CD45.1+ mice were obtained from the Swiss Immunological Mouse Repository (SwImMR). B6.129S7-Rag1^tm1Mom^/J (Jackson laboratory Stock No: 002216) mice were obtained from SwImMR. All animal work was done in accordance with the federal and cantonal laws of Switzerland. The Animal Research Commission of the Canton of Basel-Stadt, Switzerland, approved the animal research protocols.

### Adoptive T cell transfers and LCMV immunization

Edited SMARTA^+^ CD45.1^+^ CD4^+^ T cells were transferred into C57BL/6 CD45.2^+^ recipient mice by intravenous tail injections immediately after the GFP^+^ cell sorting. Mice were infected intraperitoneally with LCMV Armstrong (2*10^5^ plaque forming units) 5 days after initial T cells transfer. Mice were euthanized 3, 5 or 7 days after virus administration.

### Flow cytometry and antibodies

Cells were stained and then acquired on a BD Fortessa (BD Biosciences) and analyzed with FlowJo software (Tree Star). Surface phenotype staining was done with the following fluorochrome-conjugated mAbs: anti-CD90.2 (clone 53-2.1), anti-CD90.1 (clone OX7), anti-CD45.2 (clone 104), anti-CD45.1 (clone A20), (all eBioscience), anti-CD4 (clone RM4-5), anti-CD25 (clone PC61, both Biolegend), anti-ICOS (clone 7E.17G9), anti-PD-1 (clone RMP1-14), anti-CXCR5 (clone L138D7) (all from Biolegend) The expression of Foxp3 (clone FJK-16s) (eBioscience) was determined by intracellular staining performed according to the manufacturers’ protocols. Prior to staining of the surface antibodies cells were stained for live/dead discrimination with Zombie UV dye (Biolegend).

### Design of sgRNA

DNA sequences of all sgRNAs, primers and HDR templates used in this paper are listed as 5’-3’ sequences in the Supplementary information. sgRNAs were designed using the CRISPRtool (http://crispr.mit.edu) and sgRNA Scorer 1.0 (https://crispr.med.harvard.edu). The sgRNA sequences with their respective scores are listed in Table S1. For CD45 epitope mapping two sgRNAs were designed per candidate region. Results obtained with the ones closest to the SNP of interest are shown in the main figures. However, all 6 tested sgRNAs cut efficiently and region R1 switched epitopes with both sgRNAs (data not shown). The cut-to-mutation difference did not play a role.

### Cloning of sgRNAs into px458 plasmid

pSpCas9(BB)-2A-GFP (PX458) was a gift from Feng Zhang (Addgene plasmid # 48138). Cloning into px458 was modified from Ref (11). The px458 plasmid was digested with BbsI for 1.5h at 37°C followed by heat inactivation for 20 min at 65°C. The digested plasmid was gel-purified using the Nucleospin gel and PCR clean-up purification kit according to the manufacturer’s recommendations (Macherey-Nagel). The forward and reverse oligonucleotides (oligo) of each sgRNA were diluted at 100µM in H2O. To phosphorylate and anneal the oligos, 2µl of each oligo were mixed with T4 ligation buffer and T4 PNK to a final volume of 20µl and incubated for 30’ at 37°C (phosphorylation) followed by 5’ at 95°C and then ramping down the temperature to 20°C at -1°C/min (annealing). The annealed and phosphorylated oligos were diluted 1:200 in H2O. Ligation reactions for each sgRNA were performed by mixing 100ng of the digested and purified px458 plasmid with 2µl of the diluted phosphorylated and annealed oligos, T4 ligation buffer and T4 ligase in a final volume of 20µl. Ligation was carried out for 1h at 22°C. Bacterial transformation was performed by mixing 5µl of the ligation reaction with 50µl ice-cold chemically competent JM109 bacteria. The mixture was incubated on ice for 30 min, followed by a heat-shock at 42°C for 30’ and a subsequent 2’ incubation on ice. Then, 200µl of SOC medium (Sigma) was added and bacteria were grown for 1h at 37°C. All the transformation reaction was plated on LB plates containing 50µg/ml ampicillin. The plates were incubated overnight at 37°C. Colonies were checked for correct insertion of the sgRNA by PCR colony screening followed by sequencing. Plasmids are available from Addgene.org (Addgene plasmid numbers 82670, 82672 -82675, 82677, 102910).

### PCR colony screening for cloning into Addgene plasmid px458

Bacteria from 2 colonies per plate were picked with a pipette tip and mixed in PCR tubes with H2O, REDTaq® ReadyMix^TM^ PCR Reaction Mix (Sigma) and specific primers (forward primer GAGGGCCTATTTCCCATGATTCC, reverse primer TCTTCTCGAAGACCCGGTG). PCR was performed using an annealing temperature of 64.9°C and 35 cycles. Positive colonies (with sgRNA insertion) will display no PCR amplicon whereas negative colonies will show a 264bp amplicon.

### Plasmid sequencing

Two colonies were picked from each LB plate using a pipette tip and inoculated into a 5 ml culture of LB medium supplemented with 50µg/ml ampicillin. The cultures were grown overnight at 37°C. Plasmid DNA from the culture was isolated by GenElute Plasmid Miniprep kit (Sigma) following the manufacturer’ recommendations. Correct insertion of the sgRNA was verified by sequencing the plasmid DNA using a U6-forward primer (ACTATCATATGCTTACCGTAAC).

### HDR repair templates

DNA repair templates were designed as homologous genomic DNA sequences flanking the sgRNA binding sites. Unless noted otherwise the sgRNAs were centered as much as possible with respect to the repair templates resulting in symmetric arms of homology. Silent mutations (i.e. not altering the amino acid sequence) were introduced into the PAM sequences. Short ssDNA templates were purchased from IDT. Lyophilized ssDNA oligos were reconstituted to 10µM in ddH2O (for specific sequences see Table S2). dsDNA templates for CD90.1, CD45.1 and Foxp3 (180bp, 1kb, 2kb and/or 4kb) were purchased from Genscript as synthetic DNA cloned into pUC57 (for specific sequences see Table S3). Maxi preps (Sigma) were prepared for each of the plasmids prior to the use in the experiments. For all HDR experiments circular HDR template plasmids were used since we obtained better results compared to the use of linearized plasmids (data not shown). Plasmids containing HDR templates are available from Addgene.org (Addgene plasmid numbers 82661-82665, 82667, 104988).

### Small molecules

The following NHEJ inhibitors were used to enhance HDR: vanillin (12) reconstituted in H2O, 300µM final concentration (Sigma cat#V1104); SCR7-X in DMSO, 1µM final (Xcess Biosciences cat#M60082). Since we purchased SCR7-X from Xcess Biosciences we refer to this compound as “SCR7-X” as recently suggested (13). Rucaparib/AG-014699/PF-01367338, in DMSO, 1µM final (Selleckchem cat#S1098); veliparib/ABT-888 in DMSO, 5µM final (Selleckchem cat#S1004); RS-1 (14) in DMSO, 7.5µM final (MerckMillipore cat# 553510); RS-1 in DMSO, 7.5µM final, (Sigma cat#R9782); Luminespib/AUY-922/NVP-AUY922 in DMSO, 1µM final (Selleckchem cat#S1069); L-755,507 in DMSO, 5µM final (Tocris cat#2197); vanillin derivatives (12) 6-nitroveratraldehyde in DMSO, 3µM final (Maybridge cat#11427047), 4,5-dimethoxy-3-iodobenzaldehyde in DMSO, 3µM final (Maybridge cat#11328426); 6-bromoveratraldehyde in DMSO, 3µM final (Maybridge cat#11480124).

### Genomic DNA sequencing

Genomic DNA from different sorted cell populations (e.g. CD45.2^+^/CD45.1^−^, CD45.2^+^/CD45.1^+^, CD45.2-/CD45.1^+^, and CD45.2^−^/CD45.1^−^) was extracted by incubating the cells with the extraction buffer (100mM Tris pH 8.5, 5mM Na-EDTA, 0.2% SDS, 200mM NaCl and 100µg/ml Proteinase K; all from Sigma) for 1h at 56°C. After 15’ heat inactivation of the proteinase K at 95°C, the samples were mixed with an equal volume of isopropanol and inverted several times to facilitate DNA precipitation. After a 2’ centrifugation, the supernatant was removed and the pellet washed with 70% ethanol. DNA was pelleted by centrifugation, air dried, resuspended in milliQ water and the concentration was measured with a NanoDrop device (Witec). PCR primers including BamHI (forward TAAGCAGGATCCATTCCTTAGGACCACCACCTG) and SalI (reverse TGCTTAGTCGACACACCGCGATATAAGATTTCTGC) overhangs were purchased (Microsynth) to amplify a region of 2kb for the HDR experiment where the sgRNA location was centered within the PCR product. PCRs with 2-6ng of the different genomic DNA samples were performed using Phusion polymerase (Thermo Scientific). For the 2kb fragment the optimal annealing temperature used was 68.1°C. The PCR products were loaded on a 1.5% agarose gel and the bands were purified using the Nucleospin gel and PCR clean-up purification kit according to the manufacturer’s recommendations (Macherey-Nagel). The purified PCR products (160ng) were digested with BamHI and SalI using BamHI buffer for 1.5h at 37°C. The digested PCR products were loaded on a 1.5% agarose gel and the bands were purified using the Nucleospin gel and PCR clean-up purification kit according to the manufacturer’s recommendations. 90ng of the digested and purified 2kb PCR amplicons were ligated for 1h at 22°C with 50 or 100ng pGEM3Z plasmid which had been BamHI/SalI digested and purified (Promega). Transformation was performed by mixing 10µl of the ligation reaction with 50µl ice-cold chemically competent JM109 bacteria (purchased from Promega or made using the RbCl protocol http://openwetware.org/wiki/RbCl_competent_cell). The mixture was incubated on ice for 30’, followed by a heat-shock at 42°C for 30’’ and a subsequent 2’ incubation on ice. Then, 200µl of SOC medium (Sigma) was added and bacteria were grown for 1h at 37°C. All the transformation reaction was plated on LB plates containing 50µg/ml ampicillin, 0.1mM IPTG (Promega) and 35µg/ml x-Gal (Promega). The plates were incubated overnight at 37°C. From each plate 12 (22 for HDR) white colonies were picked using a pipette tip and inoculated into a 5 ml culture of LB medium supplemented with 50µg/ml ampicillin. The cultures were grown overnight at 37°C. Plasmid DNA from the culture was isolated by GenElute Plasmid Miniprep kit (Sigma) following the manufacturer’s recommendations. DNA was sent for sequencing using the T7, SP6 and an internal primer (GAGAAAGCAACCTCCGGTGT) for the 2kb fragments. Sequences were analyzed using Lasergene (DNASTAR Inc.)

### Cas9 RNP assembly and transfection

The delivery of a Cas9 RNP complex, containing an Alt-R CRISPR crRNA and Atto 550 labeled tracrRNA (both from IDT) and a Cas9 nuclease (from QB3 MacroLab, UC Berkeley), into primary mouse T cells or EL-4 cells using the Neon® Transfection System (Invitrogen) were adapted from a protocol provided by IDT (https://eu.idtdna.com/pages/docs/default-source/CRISPR/idt_protocol_nep-of-jurkat-rnp-rt_crs-10061-prv2-1.pdf?sfvrsn=20). In brief, the RNA oligos (crRNA and tracrRNA) were resuspended in Nuclease-Free IDTE Buffer at final concentrations of 200μM each. The two RNA oligos were mixed in equimolar concentrations to a final complex concentration of 44μM. The complexes were heated at 95º C for 5 min and then cooled down to room temperature. The 36µM Cas9 protein was pre-mixed slowly with the crRNA:tracrRNA complex and incubated at room temperature for 10– 20 min before the transfection. Fresh crRNA:tracrRNA complexes were made for each experiment as per IDT recommendations.

EL-4 cells were transfected with RNPs using the Neon® Transfection System (Invitrogen) at the following conditions: voltage (1380V), width (50ms), pulses (1) 100µl tip, buffer R (for RNPs). Primary T cells were transfected with RNPs using the Neon® Transfection System (Invitrogen) at the following conditions: voltage (1550V), width (10ms), pulses (3) 100µl tip, buffer R (for RNPs).

## Results

### Efficient plasmid-based gene ablation in a murine T cell line

We set out to develop a plasmid-based approach for CRISPR/Cas editing in primary T cells. Based on a successful T cell electroporation protocol (15) we optimized experimental conditions for murine EL-4 cells using a commonly available plasmid expressing a sgRNA, SpCas9 and GFP (11), (Fig. 1A). One day after electroporation we purified the successfully electroporated cells based on GFP expression. We quantified the efficiency of gene editing in single cells for genes encoding cell surface proteins using flow cytometry (Fig. 1B). After systematic optimization of experimental conditions taking into account electroporation parameters, concentration and choice of plasmid (size, promoter), cell number and timing, we achieved very high deletion efficiencies for CD90.2 and CD45.2, which were lost in the vast majority of cells compared to the control conditions (Fig. 1B and C). Next we asked whether this protocol allows multiplexed gene editing by combining CD45.2 and CD90.2 gene targeting. Almost half of the cells lost CD90.2 and CD45.2 expression simultaneously, indicating homozygous deletion of both genes (Fig. 1D). Thus, we successfully established very simple conditions for high efficiency gene editing in EL-4 cells.

**FIGURE 1.**
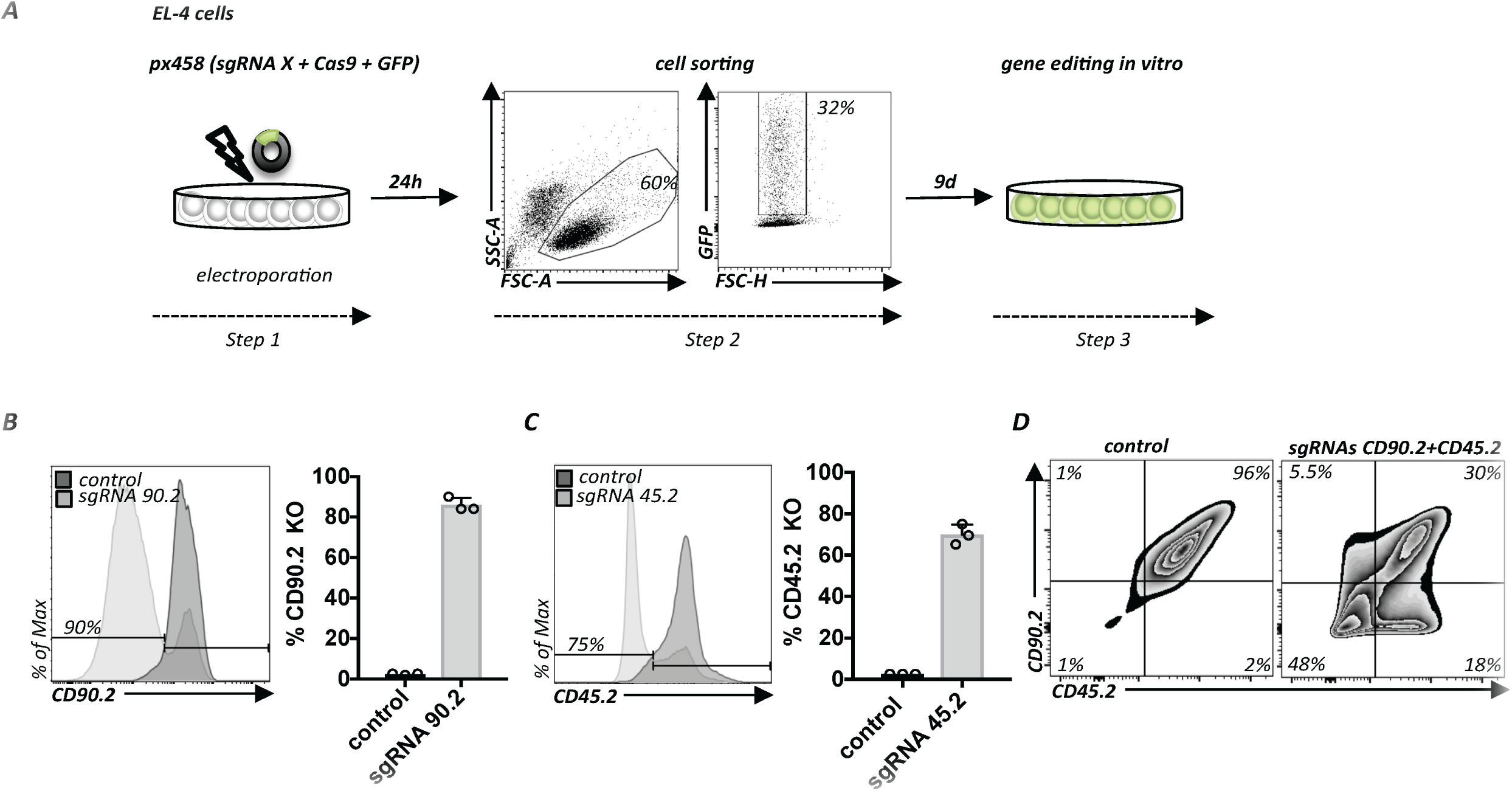
Efficient plasmid-based gene ablation in EL4 cells. (A) Protocol for plasmid-based gene editing in EL-4 cells. Electroporation of a plasmid encoding a sgRNA targeting the gene X, Cas9 and GFP (*step 1*). After 24h successfully transfected cells are purified by flow cytometry based on GFP expression (*step 2*). Subsequent cell expansion for 9 days for gene editing in vitro (*step 3*). (B) Flow cytometry of EL-4 cells transfected as in A, with a plasmid encoding a CD90.2 targeting sgRNA (sgRNA90.2) or empty vector px458 (control). Flow cytometry histograms (left panel) and quantification of multiple experiments (n=3); error bars represent standard deviation (SD) (right panel). (C) Same conditions as in B but with sgRNA45.2 or empty vector (control). Representative data from 3 experiments; error bars represent SD. (D) EL-4 cells transfected as in A but with 2 plasmids encoding 2 sgRNAs (sgRNA90.2 and sgRNA45.2). Flow cytometry of cells transfected with empty px458 vector (left panel) or cells transfected with plasmids encoding sgRNAs targeting CD90.2 and CD45.2 (right panel). Representative data from 2 experiments.

### Efficient plasmid-based gene ablation in primary T cells

Encouraged by these results we tested if the same plasmids could be used to edit primary mouse CD4^+^ T cells. To this end we added a T cell activation step with plate-bound anti-CD3 and anti-CD28 monoclonal antibodies (mAb) required for electroporation (Fig. 2A), (15). Careful titration of the concentration of mAbs used to coat the plates and timing were necessary but ultimately we found conditions allowing good viability and transfection efficiencies up to 20% (Fig. 2A). Using sgRNAs targeting CD90.2 or CD45.2 we found 60-90% deletion efficiencies among purified GFP^+^ cells (Fig. 2B and C). A second reactivation step with half the concentration of mAb was necessary for efficient editing. Remarkably, despite the notion that DNA-based CRISPR/Cas editing does not work in T cells, the gene ablation efficiencies are equal to alternative protocols using in vitro transcribed sgRNA/Cas9 mRNA or RNPs (2), (3) and even to the protocol using viral transduction of transgenic Cas9 expressing T cells (6). Next we combined the plasmids targeting CD90.2 and CD45.2. Similar to the efficiency in EL-4 cells, we found that more than half of the GFP expressing cells lost both surface proteins.

**FIGURE 2.**
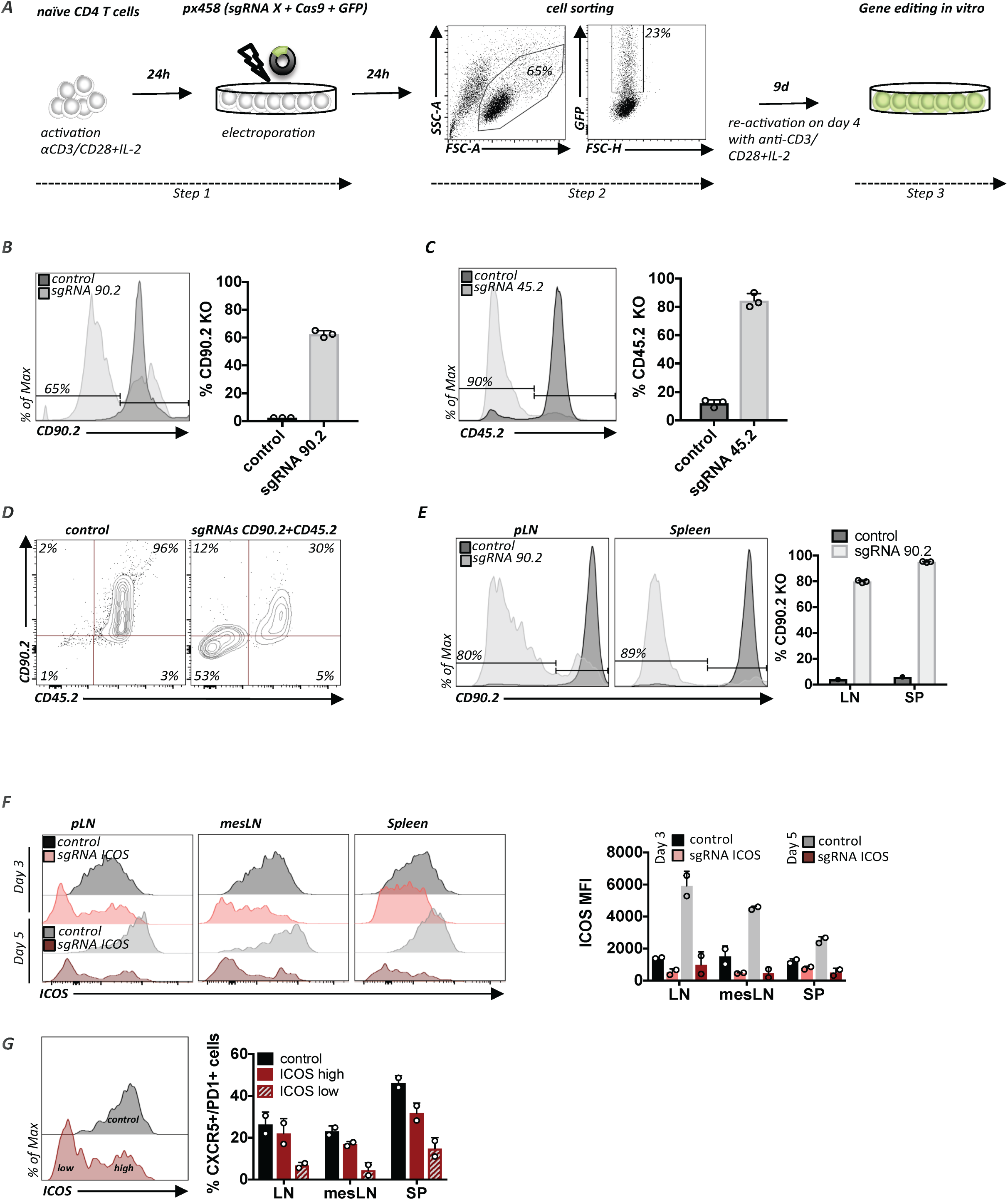
Efficient plasmid-based gene ablation in primary T cells. (A) Protocol for plasmid-based gene editing in primary CD4^+^ T cells. Activation of T cells prior to the transfection step with px458 plasmid (*step 1*). 24h later successfully transfected cells are purified based on GFP expression (*step 2*) and expanded for 9 days in vitro as shown (*step 3*). (B) Primary T cells transfected as in A, with a plasmid encoding a sgRNA90.2 or empty vector (control). Flow cytometry histograms (left panel) and quantification of 3 experiments; error bars represent SD (right panel). (C) Same conditions as in B but with sgRNA45.2 or empty vector (control). Representative data from 3 experiments; error bars represent SD. (D) Primary T cells were transfected as in A but with 2 plasmids encoding 2 sgRNAs (sgRNA90.2 and sgRNA45.2) simultaneously. Flow cytometry of cells transfected with empty px458 vector (left panel) or cells transfected with plasmids encoding sgRNAs targeting CD90.2 and CD45.2 (right panel). (E) Primary CD4^+^ T cells transfected as in A, with sgRNA90.2 or empty vector (control). Flow cytometry histograms for CD90.2 expression compared to the control on live CD4^+^ T cells in LN and SP (left panel) and quantification of multiple recipients (right panel). Representative data from 3 experiments; error bars represent SD. (F) Primary CD4^+^ T cells transfected as in A, except that CD4^+^ total T cells were used as initial population, with sgRNAICOS or empty vector (control). Flow cytometry histograms for ICOS on live CD45.1^+^ donor T cells compared to the controls in LN, mesLN and SP (left panel) at different time points and quantification of multiple recipients for ICOS MFI (right panel). (G) Demonstrates % CXCR5 and PD-1 double positive cells in LN, mesLN and SP within ICOS low and ICOS high populations (day 5) compared to the non edited control. Representative data from 2 experiments; error bars represent SD.

As mentioned previously, a powerful aspect of studying murine T cells is the possibility of adoptively transferring (AT) T cells to immunologically matched recipients. Therefore, we wondered if the edited T cells functioned in vivo. To investigate this, we adoptively transferred T cells electroporated with sgRNA for CD90.2 into lymphodeficient RAG KO mice immediately after GFP sorting. Cells were harvested from lymph nodes (LN) and spleen (SP) 10 days after AT. We observed deletion of CD90.2 in up to 80% of CD4^+^ T cells recovered from the LN and in up to 95% of CD4^+^ T cells recovered from the SP (Fig. 2 E). Thus, overall, deletion efficiencies were comparable to the ones observed in vitro. The recovered cells were viable and had expanded substantially (data not shown). Furthermore, we wanted to examine whether CRISPR/Cas9 mediated editing could be used to study gene function in primary mouse CD4^+^ T cells during a viral infection that induces T follicular helper (T_FH_) cell differentiation. We optimized experimental conditions based on a previously established protocol for acute infection with lymphocytic choriomeningitis virus (LCMV) (16). We electroporated CD4^+^ T cells bearing an LCMV-specific transgenic T cell receptor (SMARTA) with plasmid encoding a sgRNA for ICOS, a marker that is highly expressed by, and required for, T_FH_ differentiation (17). Sorted GFP^+^ T cells were transferred immediately to C57BL/6N mice. Transferred T cells were given 5 days to first migrate and then rest in host mice. Thereafter, host mice were infected with LCMV to induce T_FH_ cell differentiation. Transferred cells were harvested from LN, mesenteric LN (mesLN) and SP at 3, 5 and 7 days post virus administration. Compared to control cells we observed a progressive decrease in ICOS MFI of SMARTA^+^ CD4^+^ T cells recovered from all organs at all examined time points, with a near complete absence of ICOS expression on day 7 (Fig. 2F and Suppl. Fig. 1A). Importantly, cells that had lost ICOS expression (ICOS^low^) due to gene editing demonstrated impaired T_FH_ differentiation based on expression of the T_FH_ markers CXCR5 and PD1 compared to the cells that maintained high ICOS (ICOS^high^) expression (Fig. 2G and Suppl. Fig. 1B). Overall, this demonstrates successful gene editing and confirms the importance of ICOS signals for T_FH_ differentiation (17).

Thus, this plasmid-based approach enables efficient gene ablation in primary T cells and can be used to study T cell biology and gene function after adoptive transfer in vivo, in lymphophenic hosts but also during viral infection of lymphoreplete hosts.

### Efficient introduction of targeted point mutations in EL-4 and primary T cells

Gene editing-induced DNA double strand breaks (DSBs) are mostly repaired by non-homologous end joining (NHEJ) that results in random indels. In contrast, double strand break (DSB) repair by HDR allows controlled genome editing and is therefore desirable for more sophisticated experimental questions as well as for clinical applications. Unfortunately, as HDR occurs much more rarely, it remains challenging to establish efficient HDR protocols (18). The absence of suitable assays to readily quantify HDR events hinders improvement of HDR efficiencies in cells in general and particularly in primary cells. In order to allow rapid assessment of HDR efficiencies in T cells we designed a novel assay (Suppl. Fig. 2A-C). Two naturally occurring alleles of murine CD90 (CD90.1 and CD90.2) differ by a single nucleotide (nt) resulting in a single amino acid (aa) difference (CD90.1: arginine (Arg); CD90.2 glutamine (Gln)) (Suppl. Fig. 2B) that can be distinguished by two monoclonal antibodies (mAb), (19). We hypothesized that successful DNA editing from one allelic variant to the other could be quantified using the two allele specific mAbs (Suppl. Fig. 2B and C). To establish the allele-switching assay (ASA) we tested if we could switch the CD90.2 allele to CD90.1 in EL-4 cells by cutting with sgRNACD90.2 and providing a 180 bp ssDNA template encoding CD90.1 and a silent PAM mutation (Fig 3A left panel). Given the low HDR efficiency we observed in initial experiments (<1% CD90.1^+^ cells) we sought for ways to increase it. As it was previously demonstrated that interfering with the DNA repair pathways could increase HDR efficiencies (20), we compared several small molecules known to interfere with the NHEJ pathway, or which directly enhance HDR, to find the best HDR enhancing strategy for T cells. Along with SCR7-X, the DNA-PK inhibitor vanillin, and the PARP1 inhibitor rucaparib yielded the strongest increase in HDR frequency (Fig. 3A, right panels). Other compounds, such as veliparib, L75507 (21), luminespib, RS-1 (14) and the vanillin derivatives A14415, A1359 and L17452 (12) increased HDR less or were toxic (data not shown). Since vanillin resulted in the strongest increase in HDR and, additionally, was the only water-soluble compound we focused on vanillin for subsequent experiments. Thus, we demonstrate that allele switching of an endogenous gene can be used to quantify HDR as well as NHEJ in a population of cells with single cell resolution.

**FIGURE 3.**
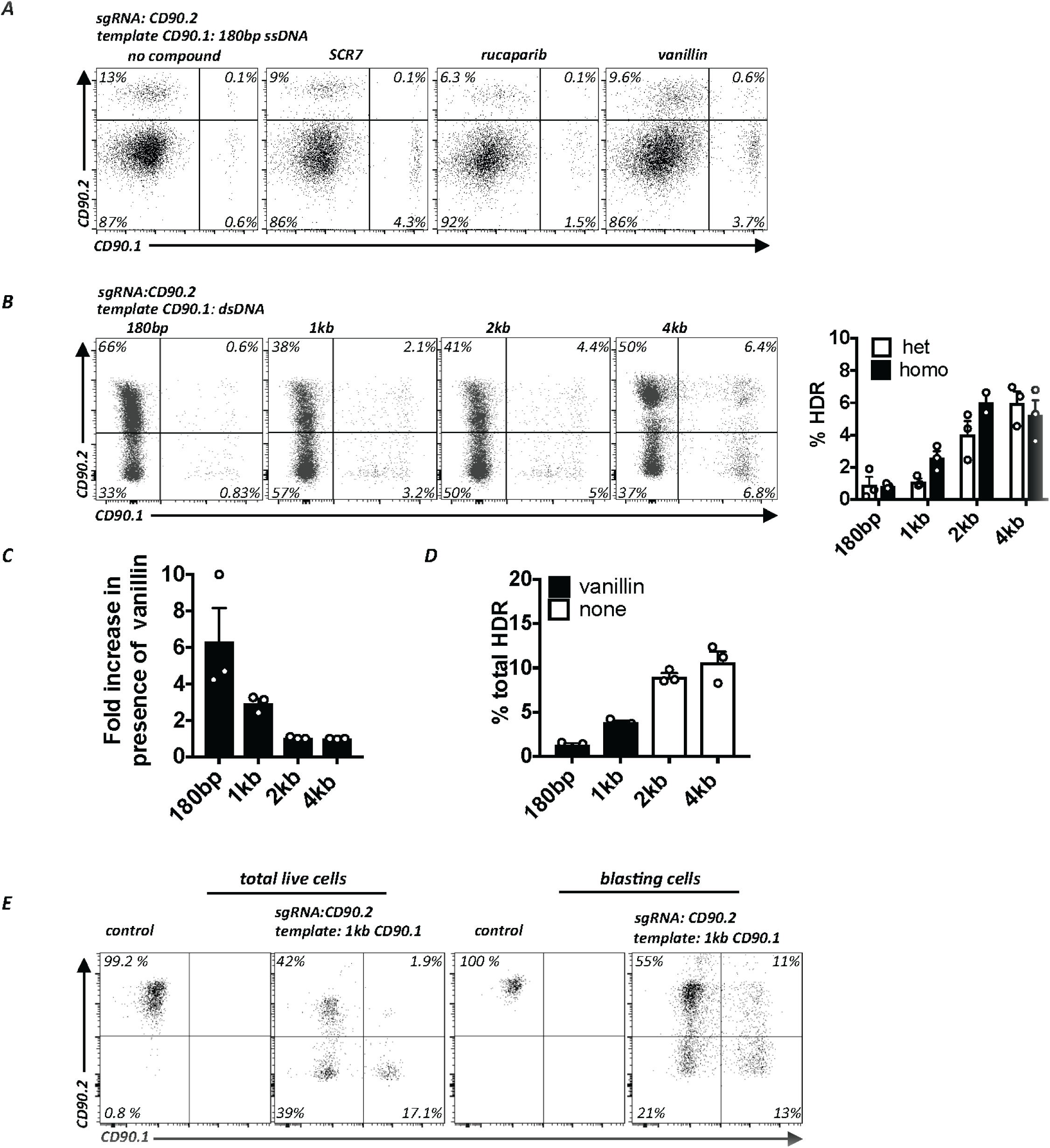
Targeted introduction of point mutations in primary T cells. (A) EL-4 cells co-electroporated with a plasmid encoding a sgRNACD90.2 and a 180bp CD90.1 ssDNA template. Flow cytometry for CD90.2 and CD90.1 expression in untreated (left panel) and treated samples (right panels). Representative data from 3 experiments. (B) EL-4 cells co-electroporated with plasmid sgRNACD90.2 and a circular plasmid including a CD90.1 dsDNA template of various lengths (180bp, 1kb, 2kb, 4kb). Flow cytometry for CD90.2 and CD90.1. Representative flow cytometry plots (left panel) and quantification of multiple experiments of the average frequency of cells that underwent HDR (heterozygous and homozygous) (right panel). Representative data from 3 experiments; error bars represent SD. (C) Quantification of the effect of vanillin on the relative enrichment of HDR frequency (fold change) as a function of dsDNA template length. Experiment as in B. Fold increase of HDR frequency of cells treated with vanillin relative to absence of vanillin for each template. Representative data from 3 experiments; error bars represent SD. (D) Long templates without NHEJ inhibitor result in higher HDR frequency than short templates with NHEJ inhibitor. Quantification of HDR frequency obtained with short templates (180bp, 1kb) plus NHEJ inhibitor (vanillin) and long templates (2kb, 4kb) without vanillin. Experiment as in B. Representative data from 3 experiments; error bars represent SD. (E) Bead-enriched naïve CD4^+^ T cells from C57Bl6/N mice activated and electroporated with empty px458 plasmid (control), with plasmid encoding for sgRNACD90.2 and a plasmid including a 1kb CD90.1 dsDNA template. Flow cytometry for CD90.2 and CD90.1 expression. Flow cytometry plots demonstrate gating on total live cells (left panels) and blasting cells (right panels). Representative data from 2 experiments.

The next parameter we evaluated was the length of the repair template. While recent gene editing reports often used relatively short ssDNA templates (usually <200bp) the arms of homology for gene targeting in embryonic stem cells are usually much longer (several kb). Indeed, increasing the arms of homology of a circular dsDNA (plasmid) CD90.1 HDR template correlated positively with HDR efficiency (Fig. 3B). The largest increase was found between 1kb and 2kb total homology (Fig. 3B). Notably, the HDR enhancing effect of vanillin was more pronounced for shorter templates (180bp, 1kb) than for the long (2kb, 4kb) templates (Fig. 3C). Therefore we wondered if a long template without NHEJ inhibition could yield a comparable HDR frequency than shorter templates with NHEJ inhibitors. A direct comparison showed that 2kb and 4kb templates without vanillin resulted in much higher HDR frequencies than the 180bp and the 1kb template in the presence of vanillin (Fig. 3D). Thus, despite the notion in the field that ssDNA templates yield higher HDR efficiencies it is worth considering long arms of homology to increase HDR efficiency. Importantly, the optimized conditions also yielded high HDR frequencies in primary mouse CD4^+^ T cells where around 20% of the total cells had switched one or both alleles (Fig. 3E, left panels). Interestingly, we noticed even higher HDR frequencies in large, blasting cells in which 25 % had undergone HDR (Fig. 3E, right panels). Thus, plasmids with long arms of homology are much more efficient to introduce point mutations to primary T cells than short ssDNA templates.

Given the many reports attributing low toxicity and high editing efficiencies to Cas9 RNPs in various cell types (NHEJ and HDR) we compared RNP performance to the results obtained with plasmids. Electroporation of EL-4 cells with recombinant Cas9 complexed with tracrRNA/crRNA targeting the same CD90.2 sequence as the one targeted by the plasmid-encoded sgRNA led to comparable KO frequencies (Suppl. Fig. 2D, left panel and Fig. 1B). In contrast, in comparison to plasmid-mediated HDR (Fig. 3B) we observed much lower HDR efficiencies when RNP was combined with the plasmid HDR template (Suppl. Fig.2D, right panel). Similar results were obtained with primary T cells (compare Suppl. Fig. 2E and Fig. 2B, 3E). Since high HDR efficiencies were reported for RNPs combined with short ssDNA templates we compared HDR templates provided as long dsDNA, or as symmetric and asymmetric short ssDNA templates, as described before (22). Compared to the all plasmid approach, HDR efficiencies were much lower with RNPs, independent of template design (Suppl. Fig. 2F).

Foreign DNA can integrate into the genome through homology-directed mechanisms or through non-homologous or “illegitimate” insertion (23). Although rare, we tested if our protocol resulted in detectable genomic plasmid integration. GFP expression used to purify successfully transfected cells peaked 24h post transfection and rapidly decreased in the following days becoming undetectable by flow cytometry after 5-7 days, both in EL-4 and primary T cells. We followed EL-4 cells up to 30 days in vitro and primary T cells up to 16 weeks in vivo and were unable to detect any GFP in these cells. However, using two different PCR primer sets detecting Cas9 or Cas9-GFP we detected faint bands in EL-4 and primary T cells at all examined time points (data not shown). We cannot tell whether this represents persisting extrachromosomal episomal DNA or true genomic integration. Future studies are required to thoroughly asses genomic plasmid integration.

Thus, the allele-switching assay described here is a simple, rapid and cost-effective system to quantify HDR efficiency with an endogenous marker in primary cells. Various HDR enhancing small molecules and HDR templates with long arms of homology (>1kb) are important parameters to consider when optimizing HDR efficiency.

### Epitope mapping of CD45.2 and CD45.1 binding antibodies in primary T cells

To test if the optimized conditions found with the CD90 ASA are more universally applicable we turned to *Ptprc*, another gene with two naturally occurring alleles, whose products CD45.1 and CD45.2 can be discriminated by two mAbs. In contrast to CD90.1 and CD90.2 however, the precise epitope recognized by mAb anti-CD45.1 (clone A20) and mAb anti-CD45.2 (clone 104) is unknown. The genomic sequence encoding the extracellular domain of CD45.1 and CD45.2 differs by 6 nt but which epitope is being recognized as an allelic difference is unknown (Fig. 4A), (25), (26). One nt substitution is silent while the other five change the aa sequence. Therefore, we hypothesized that editing the five candidate nt substitutions individually, or as combinations directly in primary T cells, could be used to fine map the epitopes being recognized by the two known mAbs. We grouped the five candidate nt into three genomic regions (R1-R3) that could be covered by three ssDNA templates. Each HDR template encoded partial CD45.1 sequences and was matched to CD45.2 cutting sgRNAs binding as close as possible to the candidate mismatches (Fig. 4A). Using the T cell HDR protocol we found that all three sgRNAs led to efficient cuts (Fig. 4B). Exchange of a single nt within region R1 enabled binding of anti-CD45.1 mAb and prevented binding of anti-CD45.2 mAb in some cells. In contrast, editing R2 and R3 did not result in anti-CD45.1 binding (Fig. 4B). A longer repair template increased HDR efficiency and confirmed this result (Fig. 4C). Sanger sequencing of all 4 purified populations (Suppl. Fig. 3A) confirmed correct editing (Suppl. Fig. 3B). Thus, the Lys302Glu substitution is necessary and sufficient to explain reactivity of the CD45.1 epitope with mAb CD45.1 clone A20 (Suppl. Fig. 3C). These results demonstrate the feasibility of epitope mapping in primary cells, i.e. in the native context of an endogenous antigen by means of using the CRISPR/Cas9 system. Furthermore, it validates the robustness of this HDR protocol and the ASA as a rapid and versatile assay to quantify single nucleotide editing.

**FIGURE 4.**
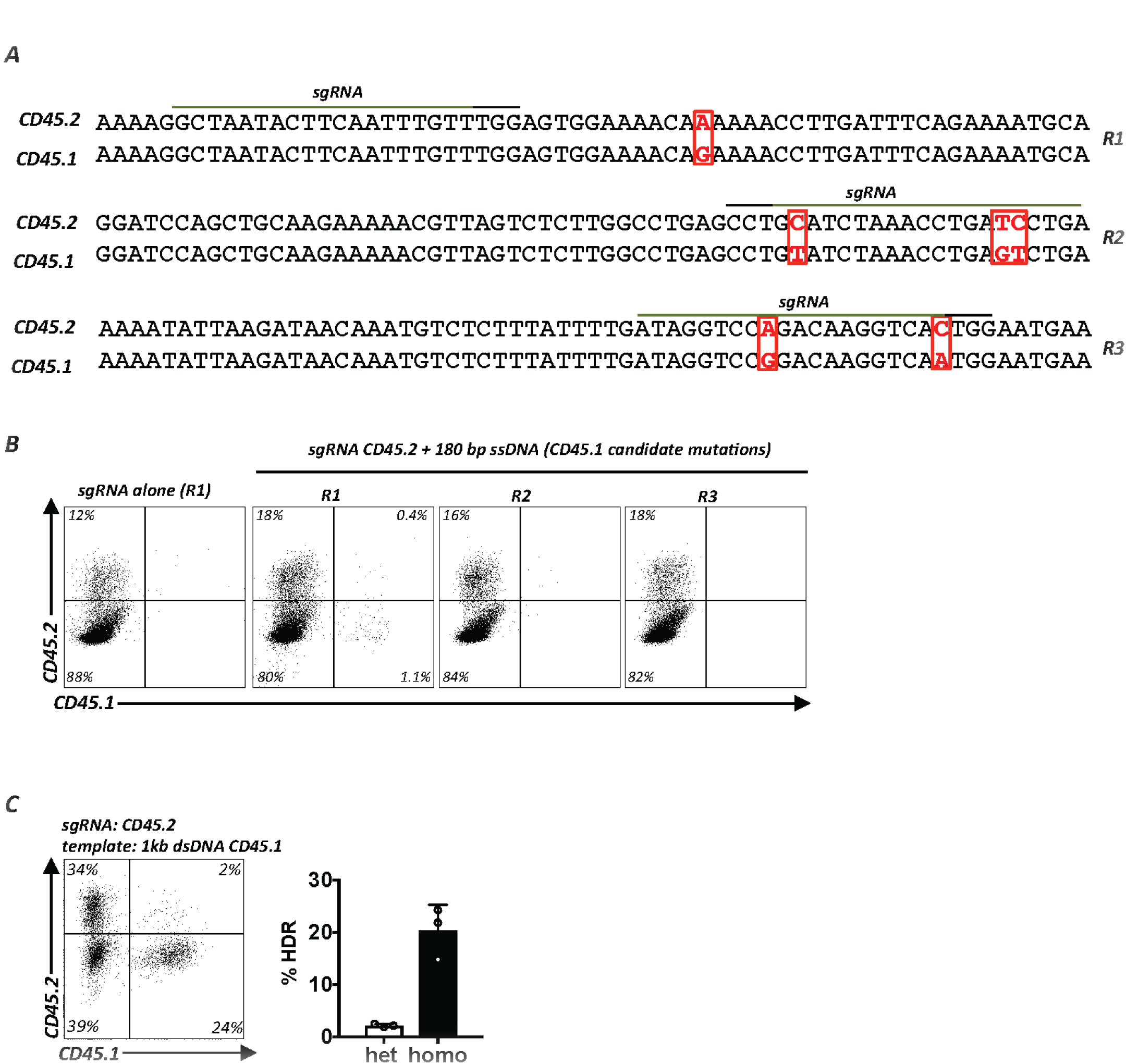
Epitope mapping of CD45.2 and CD45.1 binding antibodies in primary T cells. (A) Alignment of select regions of the genomic murine C57BL/6 CD45.1 and CD45.2 gene sequences. The extracellular domains of CD45.1 and CD45.2 differ by 6 nucleotides (indicated in red) in 3 different regions (designated R1, R2 and R3). sgRNA binding sites (green line), PAM sequence (black line). (B) High resolution gene editing-based mapping of the native CD45.1 epitope. Experimental setup as in Suppl. Fig. 2A. The three candidate regions were cut in primary CD4^+^ T cells using three different sgRNAs targeting the CD45.2 gene as close as possible to the SNP of interest (sgRNACD45.2_R1, sgRNACD45.2_R2 and sgRNACD45.2_R3) and repaired with 3 different 180bp ssDNA CD45.1 templates (R1, R2, R3). Flow cytometry for CD45.2 and CD45.1 expression. The experiment was carried out once with EL-4 cells (not shown) and once with primary CD4^+^ T cells. (C) Validation of results obtained in B using a longer 1kb CD45.1 dsDNA template. The Lys302Glu mutation is necessary and sufficient to switch CD45.2 reactivity to CD45.1 reactivity. Data are displayed as representative flow cytometry plot (left panel) and quantification of multiple experiments (right panel). Representative data from 3 experiments; error bars represent SD.

### Gene correction of Foxp3-deficient primary T cells

Next, we sought to apply the newly developed T cell editing protocol to correct a monogenic disease. The prototypic mutations causing human immunodysregulation polyendocrinopathy enteropathy X-linked (IPEX) syndrome are mutations in the *FOXP3* gene, which encodes a transcription factor critical for T regulatory cell (Treg) function, and maintenance of immune regulation (27), (28). Mutations in murine *Foxp3* lead to a very similar syndrome termed scurfy (27). A 2bp insertion in *Foxp3* exon 8 results in a frameshift leading to the scurfy phenotype (28). Affected mice die within weeks after birth due to multi-organ failure caused by a complete breakdown of immune tolerance resulting in uncontrolled activation of the immune system, tissue infiltration and immune-mediated destruction of multiple organs (29). Foxp3-deficient mice with a genetically marked *Foxp3* locus contain Treg “wanna-bes”, indicating that cells destined to become Foxp3^+^ cells are actively transcribing the *Foxp3* locus and are present in scurfy mice, but due to the absence of Foxp3, they cannot be identified as Treg and they lack suppressive function (30), (31). Thus, we hypothesized that gene correction of Foxp3 mutated T cells should lead to restoration of Foxp3 protein expression, a prerequisite for Treg function.

To test our hypothesis we used T cells from gene targeted mice that bear a Foxp3^K276X^ mutation that abolishes Foxp3 protein expression (“Foxp3 KO”) and recapitulates a known human IPEX disease-causing Foxp3 mutation (27), (31) (Fig. 5A). We adjusted the HDR-based gene repair approach to T cells from diseased mice and examined the in vitro Treg differentiation potential of gene-corrected Foxp3 KO cells by providing the Foxp3 inducing signals TGFβ alone, or as combined retinoic acid (RA) and TGFβ (32), (Fig. 5B). After gene repair and stimulation with TGFβ alone, 10% of wildtype (wt) T cells became CD25^+^Foxp3^+^ while no Foxp3^+^ cells were detected in Foxp3^K276X^ CD4^+^ T cells transfected with sgRNAFoxp3^K276X^ alone. In contrast, the Foxp3 wt repair template restored Foxp3 expression in 3.5% of the cells (Fig. 5C, top panel). Exposing electroporated T cells to combined TGFβ and RA resulted in 74% Foxp3 expression in wt T cells, no detectable Foxp3 expression in Foxp3^K276X^ CD4^+^ T cells without HDR repair template, and 18% Foxp3^+^ T cells in Foxp3^K276X^ CD4^+^ T cells repaired with the wt Foxp3 HDR template (Fig. 5C, lower panel). Comparable results were obtained with scurfy cells (data not shown). Thus, the plasmid-based HDR protocol described here works to repair primary T cells from severely sick mice.

**FIGURE 5.**
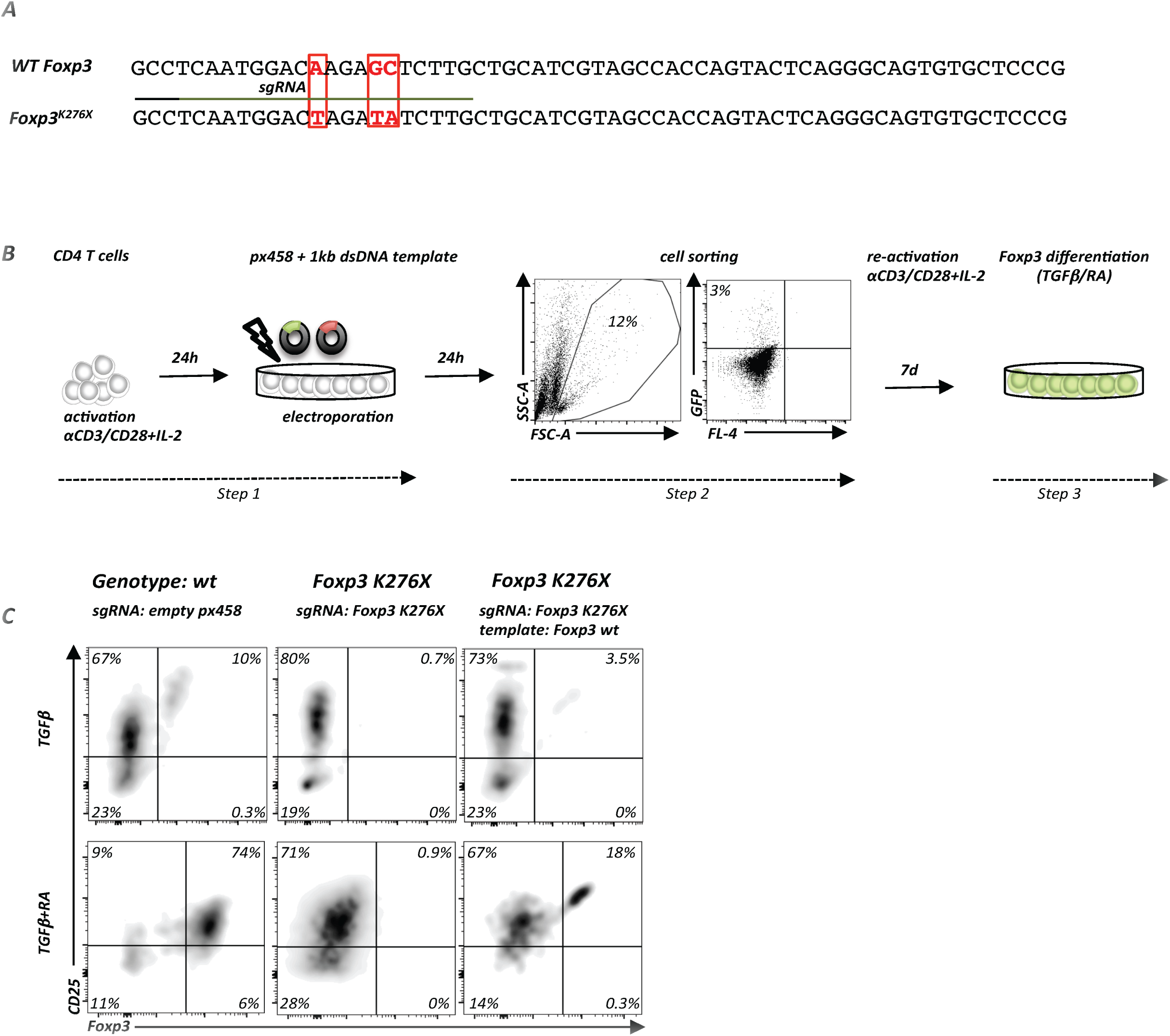
Gene correction of Foxp3 deficient cells. (A) Alignment of genomic DNA sequences of wt *Foxp3* (C57BL/6) and the *Foxp3* locus with a targeted mutation Foxp3^K276X^ that introduces a premature stop codon. sgRNA binding site (green line) and PAM sequence (black line). (B) Protocol for gene editing of total CD4^+^ T cells from Foxp3^K276X^ C57BL/6 mice. In vitro activation and electroporation (*step 1*) with plasmids encoding sgRNA targeting the Foxp3^K276X^ mutation and a circular plasmid containing a 1kb wt Foxp3 repair template. Successfully transfected cells are isolated based on GFP expression (*step 2*). Cell expansion in vitro for gene editing in presence of rhIL-2, TGFβ alone or in combination with RA and cytokine neutralizing antibodies (anti-IL-4 and anti-IFNγ) for 7 days (*step 3*). (C) Experimental setup as in B with total CD4^+^ T cells from wt control or Foxp3^K276X^ mice. Flow cytometry of CD25 and Foxp3 expression (gated on live CD4^+^ T cells). Differentiation of wt cells electroporated with empty px458 plasmid into CD4^+^Foxp3^+^CD25^+^ T cells (left panel), absence of Foxp3 differentiation in Foxp3^K276X^ cells electroporated with sgRNAFoxp3^K276X^ alone (middle panel) and restoration of Foxp3 protein expression in Foxp3^K276X^ cells electroporated with sgRNAFoxp3^K276X^ and 1kb Foxp3 dsDNA repair template (right panel). Top row: Foxp3 induction with TGFβ alone, bottom row: Foxp3 induction with TGFβ combined with RA. Representative data from 2 experiments with Foxp3^K276X^ cells.

### Two HDR events are linked in a given cell

Since we now had two unique assays at hand we wondered if the CD90 ASA and CD45 ASA could be combined to quantify multiplexed HDR in single cells. In order to address this question, we electroporated EL-4 cells with plasmids encoding sgRNAs targeting CD90.2 and CD45.2 along with repair templates for CD90.1 and CD45.1. Cutting efficiency under these conditions was slightly lower than with fewer plasmids (data not shown) but HDR for CD90 and CD45 individual alleles was very efficient. We then sought to determine if two HDR events at two separate loci in the same cell are independent from each other or linked. We found a 2-fold enrichment of cells switching CD45.2 to CD45.1 in cells that had switched CD90.2 to CD90.1 compared to cells that remained CD90.1^-^ (Fig. 6A). Importantly, a third of the CD90.2+/CD90.1^+^ heterozygous cells were also heterozygous for CD45.2^+^/CD45.1^+^ (Fig. 6B). Similarly, the highest relative frequency of homozygous CD45.1^+^ cells was found among cells that were also homozygous for CD90.1^+^ (Fig. 6B). Thus, DNA repair by the HDR pathway is more likely to occur in cells that concurrently repair a second DNA break by HDR.

**FIGURE 6.**
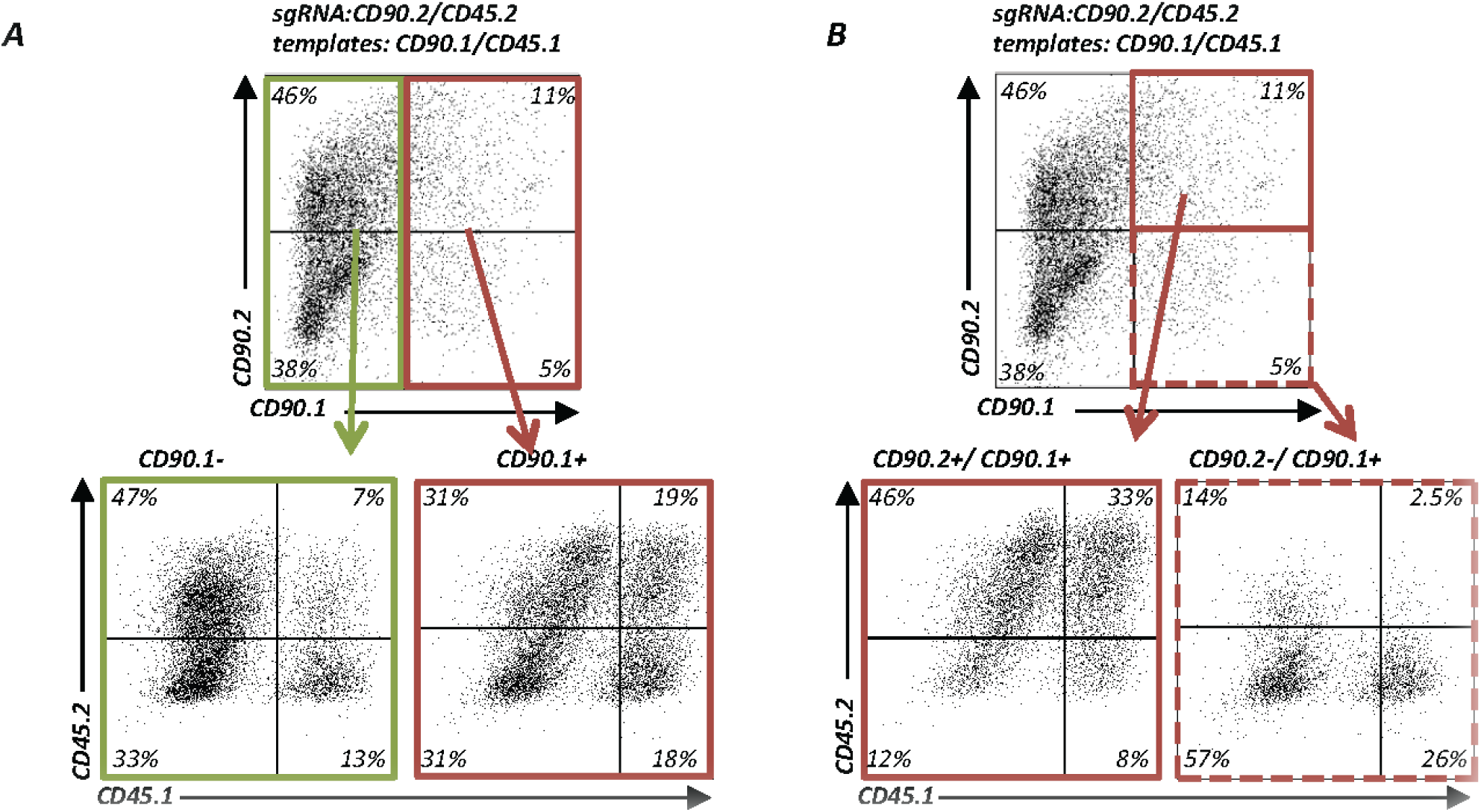
Enrichment of HDR-edited cells through monitoring of allele switching of a surrogate cell surface marker. (A) Enrichment of HDR-edited cells using allele switching of a surrogate cell surface marker. EL-4 cells electroporated with plasmids encoding sgRNACD90.2 and sgRNACD45.2_R1 and 2kb dsDNA templates (CD90.1 and CD45.1) for multiplexed HDR. Flow cytometry for CD90.2, CD90.1, CD45.2 and CD45.1 expression. Top panel: pre-gating on CD90.1^−^ (green) and CD90.1^+^ (red) i.e. allele switched cells demonstrates that HDR events at a second locus (*Ptprc*) are linked within the same cell. CD45 allele switched cells (lower panels) are more frequent in cells, which also switched the CD90 allele. Representative data from two experiments. (B) Selection of zygosity of HDR-edited cells. Experimental data as in A. Top panel: pre-gating on heterozygous CD90.1+/CD90.2^+^ cells (solid red line) enriches CD45.1^+^/CD45.2^+^ heterozygous cells (left bottom panel). Pre-gating on homozygous CD90.1+/CD90.1^+^ cells (top panel, dotted red line) enriches homozygous CD45.1^+^/CD45.1^+^ cells (bottom panel).

### Allele switching of a surrogate surface receptor enriches gene-repaired cells

Finally, we wondered if linked HDR could be exploited to enrich correctly repaired cells using multiplexed HDR. In a clinical context it will be desirable to select correctly edited cells before transfusion to patients. However, identification of these cells (without killing them) is difficult if the repaired gene is not expressed at the cell surface. Since Foxp3 is an intracellular transcription factor we sought to repair primary Foxp3 KO CD4^+^ T cells and combine the repair process with CD45.2 to CD45.1 allele switching as a surrogate cell surface marker to monitor allele switching. As described in Fig. 5, total CD4^+^ T cells were isolated from Foxp3^K276X^ mice and electroporated with four plasmids to cut CD45.2 and mutant Foxp3 and repair them with CD45.1 and wt Foxp3 templates. Thereafter, cells were exposed to TGFβ to induce Foxp3 expression. Indeed, CD25^+^Foxp3^+^ cells were substantially enriched among CD45.1^+^ cells (16%) compared to CD45.1^-^ cells (0.1%) (Fig. 7A).

**FIGURE 7.**
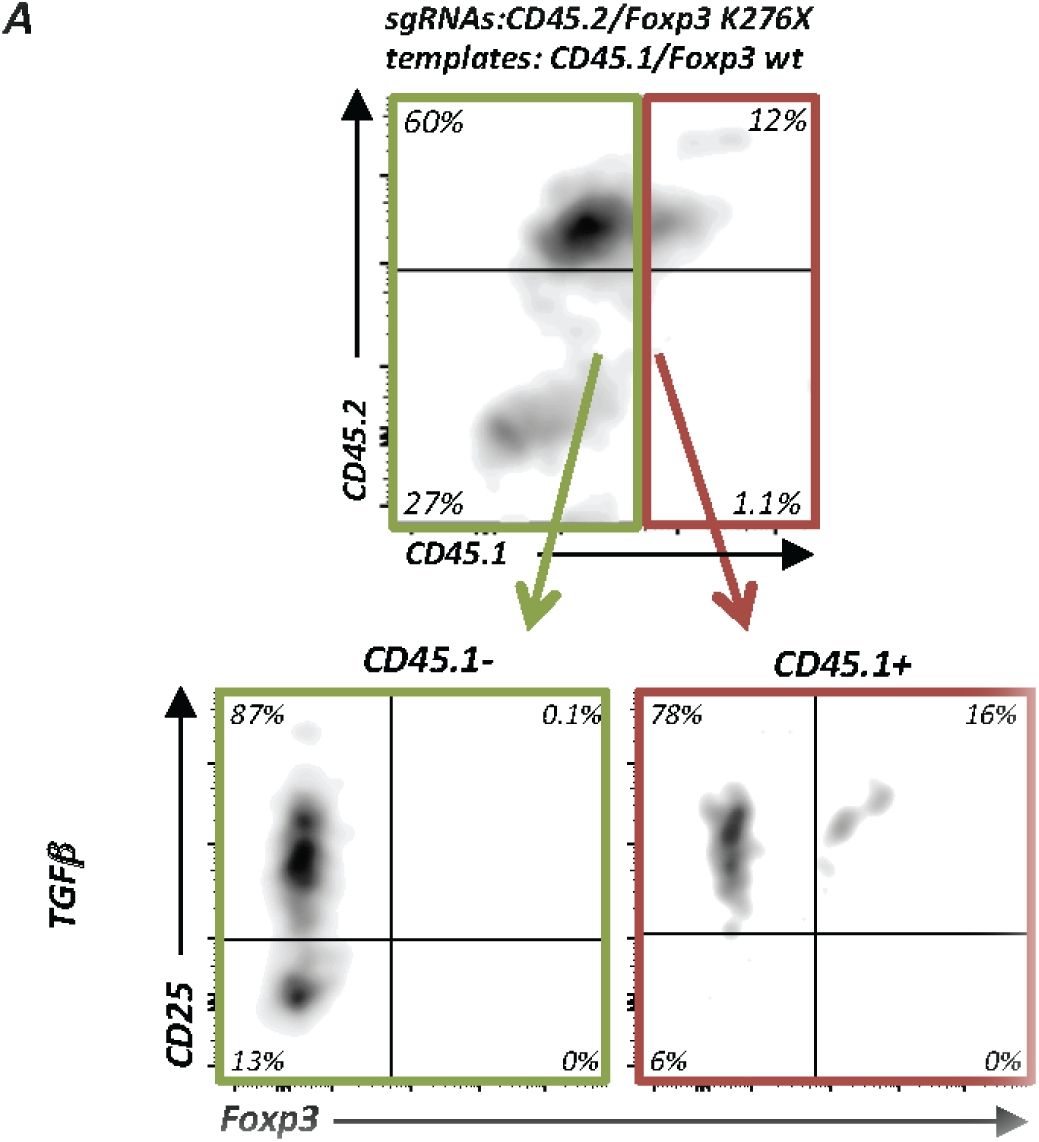
Enrichment of Foxp3 repaired cells through monitoring of allele switching of a surrogate cell surface marker. (A) Enrichment of gene-repaired Foxp3 expressing cells using multiplexed CD45 allele switching as a surrogate marker. Experimental setup as in Figure 5B but simultaneous electroporation of plasmids encoding 2 sgRNAs (sgRNAFoxp3^K276X^ and sgRNACD45.2_R1) and two 1kb dsDNA templates (Foxp3 wt and CD45.1). Flow cytometry of CD45.2, CD45.1, CD25 and Foxp3 (gated on live CD4^+^ cells). Top panel: pre-gating on CD45.1^−^ cells (green line) and CD45.1^+^ cells (red line). Bottom panel: Enrichment of CD25^+^Foxp3^+^ cells in allele switched CD45.1^+^ cells. Representative data from 2 experiments.

Compared to Foxp3 repair without allele switching, which under these conditions (TGFβ alone) resulted in 3.5% Foxp3 expressing cells (Fig. 5c, upper panel), monitoring allele switching of a surrogate surface marker resulted in approximately 4- fold enrichment. In summary, we established conditions to repair *Foxp3* in primary T cells and demonstrate the applicability of multiplexing HDR to enrich gene-corrected cells. We therefore propose that assessment of a surrogate marker HDR gene editing event could be exploited to enrich and/or select for zygosity of HDR gene editing at a second gene locus of interest for which no marker is available.

## Discussion

Efficient protocols to genetically engineer T cells will likely contribute to a better understanding of the architecture of our genome and the wiring of genetic networks guiding cellular behavior, and will, therefore, ultimately lead to safer and more efficient cellular therapies. In contrast to recent results suggesting that plasmid-based genome engineering is inefficient in T cells, our results demonstrate that plasmids are well-suited, very powerful and versatile vectors to edit a T cell’s genome. Plasmids are commonly available, inexpensive, transiently expressed vectors which can be used to edit the many existing genetically modified mouse models including T cell receptor transgenic mice. Importantly, the method described here does not rely on importing and intercrossing mice but can immediately be applied to cells of existing mouse models, independent on genetic backgrounds. Thus, we expect that this method will accelerate research in areas as diverse as genome biology, development, lymphocyte biology, immunology and animal models of cellular therapy. It will be important to investigate potential genomic plasmid integration and off-target mutations. Since plasmids tend to persist longer in cells than RNPs and since by PCR we detect plasmid sequences in some cells up to several weeks after electroporation, the frequency of off-target cleavage could be increased compared to the use of RNPs. For many basic research questions this may not be relevant as off-target cleavage can be controlled for by using different sgRNAs. In contrast, for potential clinical translation these questions are highly relevant and warrant thorough investigation.

Our results go beyond the field of immunology however. Although targeted HDR is often preferred over NHEJ, achieving high efficiency HDR is still challenging. The results from the allele switching assays suggest that HDR templates with long arms of homology should be revisited to increase HDR efficiency. In addition, while NHEJ inhibitors can be useful to increase HDR, in cases where NHEJ inhibitors are preferentially avoided (e.g. clinical gene editing), long arms of homology can prove effective and do not necessarily have to be virally delivered. Although we used dsDNA HDR templates it will be worthwile to investigate if increasing the length of the arms of homology of ssDNA templates leads to comparably increased HDR efficiency. Using long ssDNA templates rather than dsDNA templates might also decrease the risk for off-target genomic HDR template integration. Moreover, we provide proof-of-concept data that genome-editing can be applied for rapid and precise epitope mapping in primary cells. This approach could be particularly powerful to characterize the majority of B cell epitopes, which are discontinuous, i.e. conformational. The approach presented here is unique since the mapping was achieved in primary cells, i.e. in the native context with all endogenous posttranslational modifications. Knowing precise epitopes is particularly important for the many therapeutic antibodies but also to define the binding sites of chimeric antigen receptors (CAR) that are derived from mAbs. Since tumors can escape attacks by the highly efficient CAR-T cells, gene editing-based epitope mapping could be applied to tumor cells to investigate escape mutants or to define tumor antigens.

Finally, cell-based therapeutics constitute the next “pillar” of medicine (33). Genome modification of the cellular product can repair genetic defects before subsequent autologous transplantation and allows equipping the cell with designer features to increase safety and efficacy (33), (34). Thus, protocols to efficiently introduce precise and targeted genetic modifications in hematopoietic cells, particularly T cells, are key to the success of adoptive T cell therapy. Targeted insertion of a CAR into the endogenous T cell receptor locus is likely not only safer but also results in more efficient CAR-T cells compared to virally delivered randomly integrated CAR constructs (7), (35). To further optimize cellular products and to thoroughly investigate novel, experimental synthetic genetic networks, mouse models will remain important animal models for T cell therapy. To this end, this plasmid-based protocol enables efficient, non-viral, targeted cellular engineering to investigate new concepts for cellular therapies in immunocompetent mouse models (36). We would like to caution however, that adaptation of this protocol to human T cells, particularly for clinical use, would require a thorough investigation of potential genomic plasmid integration as well as possible other effects such as triggering of TLR9.

## Acknowledgments

We would like to thank the University of Basel, the Basel University Hospital and Department of Biomedicine (DBM) for institutional support, the lab members, Pawel Pelczar and the members of the DBM genome editing club for discussions, Marianne Dölz, Oliver Gorka, Jeffrey Bluestone, Xuyu Zhou and Tony Nguyen for critical comments on the manuscript and Regan Geissman for editorial assistance. Angelika Offinger, Ulrich Schneider and team from the DBM animal facility for animal husbandry. Ed Palmer and Carolyn King for kindly sharing mice, Georg Holländer for kindly sharing EL-4 cells, Shane Crotty and Simon Bélanger for sharing ICOS sgRNA sequences, Annaise Jauch for providing LCMV. Danny Labes, Emmanuel Traunecker and Lorenzo Raeli of the DBM flow cytometry core for support and Marco Amsler and Caroline Schwenzel for technical support.

## Supplemental Figure legends

SUPPLEMENTAL FIGURE 1. Plasmid-based ICOS ablation prevents LCMV induced T_FH_ differentiation. (A) Primary CD4^+^ T cells transfected as in Fig 2A, except that CD4^+^ total T cells were used as initial population, with sgRNAICOS or empty vector (control). Flow cytometry histograms for ICOS on live CD45.1^+^ donor T cells relative to the non-edited cells in LN, mesLN and SP (left panel) and quantification of multiple recipients for ICOS MFI (right panel). (B) Demonstrates CXCR5 and PD-1 double positive cells in LN, mesLN and SP within cells that were transfected with sgRNAICOS compared to the non-edited control. Representative data from 2 experiments; error bars represent SD.

SUPPLEMENTAL FIGURE 2. Allele switching assay to monitor and optimize HDR based editing efficiencies. (A) Protocol for plasmid-based HDR in CD4 T cells. Activation and electroporation of primary T cells (*step 1*). Purification of GFP^+^ cells by flow cytometry (*step 2*). Cell incubation for 24h with NHEJ inhibitor. Subsequent in vitro cell expansion for gene editing for 6-9 days with reactivation 4 days post sorting (*step 3*). EL-4 cells are transfected the same way, except they do not require TCR activation prior to the electroporation or on day 4 post sorting and electroporation parameters are different (see Materials & Methods). (B) Genomic CD90.1 and CD90.2 nt and aa sequences. The CGA (CD90.1)/CAA (CD90.2) SNP leading to Arg108Gln is highlighted in red. (C) Graphic representation of the experimental readout: Q1: unedited cells or cells with mutations which do not abolish protein expression, e.g. in-frame mutations Q2: cells after NHEJ Q3: edited CD90.2/CD90.1 heterozygous cells Q4: edited homozygous CD90.1 cells or cells with one KO allele and one HDR edited allele. (D) EL4 cells transfected with Cas9 RNP targeting CD90.2 alone or with 2 kb dsDNA CD90.1 DNA template. Flow cytometry for CD90.2 and CD90.1. (left panel) and quantification of 2 experiments; error bars represent SD (right panel). (E) Primary CD4^+^ T cells transfected with Cas9 RNP targeting CD90.2 alone or with 2kb dsDNA CD90.1 DNA template. Flow cytometry for CD90.2 and CD90.1. (left panel) and quantification of 2 experiments; error bars represent SD (right panel). (F) EL4 cells transfected with Cas9 RNP or px458 plasmid targeting CD90.2 alone or with CD90.1 DNA templates provided as symmetric and asymmetric 180bp ssDNA templates or as a plasmid including a 2 kb dsDNA template. Quantification of KO efficiency and HDR in presence of symmetric, asymmetric short ssDNA, or long dsDNA template between plasmid based and RNPs approach.

SUPPLEMENTAL FIGURE 3. Sanger sequencing of HDR edited populations. (A) Pre- and post-sort frequencies and purity of the cell populations isolated for DNA sequencing. (B) EL-4 cells electroporated with a plasmid encoding a sgRNACD45.2 and a circular dsDNA plasmid carrying 2kb CD45.1 as described in Suppl. Fig.2A. Cells were cultured for 9 days in vitro, then harvested and sorted by flow cytometry based on CD45.2 and CD45.1 expression in order to isolate four defined populations: CD45.2^+^/CD45.1^−^ (Q1), CD45.2^−^/CD45.1^−^ (Q2), CD45.2^+^/CD45.1^+^ (Q3) and CD45.2^−^ /CD45.1^+^ (Q4). DNA was extracted and cloned for Sanger sequencing as described in materials and methods. No indels were found at both ends of the templates for populations Q3 and Q4 (data not shown). (C) Genomic CD45.1 and CD45.2 nt and aasequences. The GAA (CD45.1) AAA (CD45.2) SNP leading to Lys302Glu are highlighted in red.

## Supplementary tables

Table S1. sgRNA design and their specific sequences.

Table S2. Single strand (ss) DNA templates for HDR repair experiments.

Table S3. Double strand (ds) DNA templates for HDR repair experiments.

## Footnotes

2 M.K. performed and analyzed all experiments in the manuscript. R.M. performed cloning of PCR products for sequencing and analyzed sequencing data, helped cloning and depositing plasmids, helped to write materials and methods and to prepare the supplementary tables; M.K. and L.T.J. designed the experiments, interpreted the data, discussed results and wrote the manuscript.

3 Data and materials availability: All plasmids used in the manuscript are deposited at addgene.org

4 Address correspondence and reprint requests to Lukas T. Jeker, MD PhD, Assistant Professor of Experimental Transplantation Immunology & Nephrology, Molecular Immune Regulation, Lab 313, Department of Biomedicine, Basel University Hospital and University of Basel, Hebelstrasse 20, CH-4031 Basel, Switzerland;

5 The online version of this article contains supplemental material.

6 Competing interests: M.K. and L.T.J. have filed provisional patent applications related to this work.

7 Abbreviations used in this article: ASA, allele switching assay; ACT, adoptive cell transfer; CAR, chimeric antigen receptor; Cas9, CRISPR associated protein 9; CRISPR, clustered regularly interspaced short palindromic repeat; crRNA, CRISPR RNA; DSB, double strand break; dsDNA, double stranded DNA, gRNA, guide RNA; HDR, homology directed repair; hHSC, human hematopoietic stem cells; IPEX, immunodysregulation polyendocrinopathy enteropathy X-linked; LCMV, lymphocytic choriomeningitis virus; NHEJ, non homologous end joining; PAM, protospacer adjacent motif; RNP, ribonucleoprotein; sgRNA, single guide RNA; SNP, single nucleotide polymorphism; ssDNA, single stranded DNA; tracrRNA, trans-activating crRNA;

